# DECODING COMPLEXITY IN BIOMOLECULAR RECOGNITION OF DNA I-MOTIFS

**DOI:** 10.1101/2023.04.19.537548

**Authors:** Kamyar Yazdani, Srinath Seshadri, Desiree Tillo, Charles Vinson, John S. Schneekloth

**Affiliations:** Chemical Biology Laboratory, National Cancer Institute, 1050 Boyle St., Frederick, MD 21702; Genome Analysis Unit National Cancer Institute, 3.7 Convent Dr., Bethesda, MD 20892; Laboratory of Metabolism Center for Cancer Research National Cancer Institute, 37 Convent Dr., Bethesda MD 20892

**Keywords:** DNA recognition, DNA structures, i-motif, fluorescent probes, high-throughput screening

## Abstract

DNA i-motifs (iMs) are non-canonical C-rich secondary structures implicated in numerous cellular processes. Though iMs exist throughout the genome, our understanding of iM recognition by proteins or small molecules is limited to a few examples. We designed a DNA microarray containing 10,976 genomic iM sequences to examine the binding profiles of four iM-binding proteins, mitoxantrone, and the iMab antibody. iMab microarray screens demonstrated that pH 6.5, 5% BSA buffer was optimal, and fluorescence was correlated with iM C-tract length. hnRNP K broadly recognizes diverse iM sequences, favoring 3-5 cytosine repeats flanked by thymine-rich loops of 1-3 nucleotides. Array binding mirrored public ChIP-Seq datasets, in which 35% of well-bound array iMs are enriched in hnRNP K peaks. In contrast, other reported iM-binding proteins had weaker binding or preferred G-quadruplex (G4) sequences instead. Mitoxantrone broadly binds both shorter iMs and G4s, consistent with an intercalation mechanism. These results suggest that hnRNP K may play a role in iM-mediated regulation of gene expression *in vivo*, whereas hnRNP A1 and ASF/SF2 are possibly more selective in their binding preferences. This powerful approach represents the most comprehensive investigation of how biomolecules selectively recognize genomic iMs to date.

## Introduction

The three-dimensional structure of the genome has a major impact on gene expression. Predominantly, the human genome consists of duplex B-form DNA though it can also adopt several non-B conformations. Included among these diverse structures are tetraplexes, which are represented by G4s as well as the lesser-studied intercalated-motif (iM). Unlike G4s, which are characterized by Hoogsteen-bonded tetrads of guanines coordinated by central potassiums, iMs are stabilized by intercalated hemi-protonated cytosine nucleobases (C:C+)^[1]^. It has been shown that G4 and iM formation is interdependent, indicating that they may act as components of a regulatory switch mechanism^[2]^. iM-forming sequences are enriched in CG islands^[1]^, regulatory regions of the genome, oncogene promoters (*c-MYC, KRAS, BCL-2*, and *VEGF*), untranslated regions (UTRs)^[3]^, centromeres^[4]^, and telomeres^[3, 5]^. Consequently, iMs are thought to be broadly involved in regulating essential biological processes such as transcription, cell cycle progression, DNA damage repair, and telomere maintenance. However, little is known about how diverse iM sequences recognize or distinguish between different binding partners. To date, high-throughput studies investigating protein or small molecule binding to genomic iMs are lacking. A broad understanding of how iMs achieve specificity in biomolecular recognition would provide crucial insights toward rationally targeting iMs, helping to clarify their biological significance, and potentially informing the development of novel therapeutics. The first iM discovered consisted of a d(TCCCCC) repeat,^[6]^ but it is now widely acknowledged that iMs form over a diverse range of sequence space. Parallel strands can intercalate in multiple different topologies and are influenced by the length of cytosine tracts (ranging from 3 to 12 nucleotides^[7]^), the presence of flanking thymine residues, the length of lateral loops, and torsional stress from associated proteins^[8]^. It has also been shown that longer iMs of d(TCCC)_*n*_ where *n* ≥ 7, are able to fold near physiological pH *in vitro*^*[9]*^. The sequence composition of loops in the iM further contribute to the relative stability of the structure. Thymidine is commonly a terminal residue, acting as a cap for the iM by similarly forming T:T base pairs^[1]^. The diversity of sequences capable of forming iMs means that many thousands of sequences throughout the human genome have the potential to fold into iM conformations. A key aspect of the iM structure is the pH sensitivity of the C:C+ basepair, which is mainly dictated by the pKa of the N3 cytosine (∼4.6)^[10]^. This acid-dependency led to questions about its propensity to form under physiological conditions. However, iMs were successfully visualized in the nucleus of live cells with a structure-specific scFv called iMab. This seminal work demonstrated that iM formation in the nucleus was cell-cycle dependent, in which iMs were most prevalent in the late G1 phase, which is characterized by increased transcription. Thus, a credible basis for the iM as a biologically-relevant regulatory structure was established, encouraging further studies^[11]^.

Many since have demonstrated that both proteins and small molecules readily recognize iM structures. Small molecule ligands that bind to iMs in oncogene promoters can also impact gene expression, where they frequently have the inverse effect of G4 stabilization. For example, the small molecules IMC-48 and IMC-76 encourage the formation of an iM or a flexible hairpin in the *BCL2* promoter, respectively. Stabilization of the *BCL2* iM significantly upregulated downstream expression, suggesting a potential oncogenic role^[12]^. Similarly, a bisacridine derivative was shown to stabilize *c-Kit* promoter iMs, attenuating oncogenic transcription and translation^[13]^. Other examples of iM-binding ligands include a ruthenium(II) flavone that binds to the *c-MYC and VEG-F* iMs^[14]^, mitoxantrone, pyridostatin, an amino-substituted pyrimidine that binds to the *HRAS* iM^[15]^, and a peptide-nucleic acid (PNA) oligomer that binds to the *KRAS* iM^[16]^. In addition, several different endogenous proteins are known to bind to iMs, where they frequently unfold the structure. For example, BmILF, hnRNP LL, and hnRNP K have all been shown to bind with high specificity to iMs found in promoter regions of oncogenes such as *KRAS* and *c-MYC*^*[12]*^. A broad understanding of iM binding profiles is necessary to understand the selectivity of interactions between iMs and synthetic ligands or endogenous proteins. Previously, we and others^[17]^ have demonstrated the advantages of using a DNA microarray-based platform to probe the binding specificities of G4-forming sequences against different proteins, antibodies, and small molecules^[18]^. This approach allows for a highly sensitive quantification of binding interactions for over thousands of G4-forming sequences in parallel and provides important insights into the binding of both endogenous and synthetic ligands.

In this study, we report the design of custom DNA microarrays with 10,976 distinct iM-forming sequences found in human promoters, centromeres, and telomeres, with varied lengths of cytosine repeats ranging from two to ten bases and loops ranging from one to ten bases (Table 1). Using a Cy5-labeled iMab antibody, we demonstrate proper iM folding on the microarray, and quantify binding events to determine the influence of cytosine tract length on iMab recognition. We also investigate the effect of pH and molecular crowding on iM folding propensity during these experiments. Select iMs from the array that bound the iMab antibody were analyzed by circular dichroism to further elucidate the influence of nucleotide sequence and buffer conditions on iM thermostability. This approach is also used to investigate four reported iM-binding proteins (hnRNP K, hnRNP A1, hnRNP LL, and ASF/SF2^[19]^) as well as the known iM-binding small molecule mitoxantrone, to further gain insights into sequence or structure-dependent binding specificities. We also compare binding data on the array to public ChIP-Seq data, confirming interactions between hnRNP K and iM-forming sequences in cells. Taken together, these results demonstrate that array-based techniques can provide a broad and accurate perspective of the binding preferences of relevant biomolecules to genomic iM structures.

**Table 1.** iM DNA microarray design. 55,000 total iMs were included in the array, in addition to 5,000 positive controls, negative controls (iM-forming sequences with all Cs mutated to T or A), G4s, and non-G4s (unstructured ssDNA). Additional negative controls added included 295 G4-forming sequences and 46 non-G4-forming sequences. A 3’ TTTTTTTT linker was added to extend all sequences from the array surface, encouraging proper folding.

| Sequence Class | Number | Replicates |
| --- | --- | --- |
| 5-10Cs | 5,102 | 5 |
| 4Cs (promoter) | 3,176 | 5 |
| 3Cs (promoter) | 2,000 | 5 |
| 2Cs (promoter) | 682 | 5 |
| Telomere | 12 | 10 |
| Centromere | 4 | 20 |
| Positive Controls | 8 | 100 |
| Negative Controls | 500 | 5 |
| G4s | 300 | 5 |
| Non-G4s | 40 | 5 |

## Results and Discussion

To design custom iM DNA microarrays, the hg38 human genome was parsed for putative iM sequences (Table 1). We utilized a consensus iM formula, C_*n*_(N_1-19_C_*n*_)_3_, where N is any nucleotide, and *n* is the number of cytosines in the iM C-tract^[7]^. Additionally, we specified a maximum iM length of 52 nucleotides. With these parameters, we first searched the genome for 5-10C iMs, identifying 5,102 putative 5-10C iM sequences. Next, we looked through promoter regions of the hg38 genome by searching the -1kb regions of Gencode v36 annotated genes in both the sense and antisense directions.

Here, we found 3,176 4C iMs, 65,129 3C iMs, and 682 2C iMs, in which we randomly selected 2,000 3C iMs for the array. 5 replicates of each sequence were added to the array. For telomere sequences, we searched the genome for the consensus telomere repeat (TAAC_3_)_5_ identifying 12 distinct telomeric regions, and 10 replicates of each sequence were added. Centromeric sequences identified previously in the literature, specifically the 2 CENP-A and B box sequence variants, were printed on the array 20 times^[1]^. Positive control sequences were added based on the argument that the most stable iMs contain first and third loops with three thymines. Thus, the consensus positive control sequence formula, C_*n*_(C_*n*_T_3_)_3_, where 3 ≤ *n* ≤ 10, was used to identify 8 positive control sequences, in which 100 replicates were incorporated^[7, 20]^. Negative control sequences were designed by randomly selecting 500 4C iMs and replacing all C-tracts with an A or T, in which both have an equal chance of being selected. These mutant sequences were printed 5 times on the array. Additional negative controls included 295 G4-forming sequences and 46 non-G4-forming sequences, in which 5 replicates were printed. A 5’ TTTTTTTT linker was added to all sequences on the array to extend the sequence from the array surface, encouraging proper folding. It has been established that an acidic solution with inert crowding is ideal for *in vitro* iM formation (Figure 1a)^[21]^. We evaluated binding at pHs 5.5, 6.5, and 7.5 and modulated bovine serum albumin (BSA) concentration from 5% to 40% (w/v) in our array hybridization buffer, using iMab to determine proper iM folding (Figure 1c-f). pH 6.5 with 5% BSA achieved the highest mean difference between the positive and negative control signal to noise ratios (SNR) upon iMab incubation and was used for subsequent microarray screens. A similar difference was observed at pH 7.5, 40% BSA, suggesting that crowding agents can play a stabilizing role under physiological conditions through negative superhelical force, such as in the case of the *c-MYC* promoter iM^[22]^. Aside from pH, it has been shown that the stability of an iM is positively correlated with the number of consecutive cytosines in their C-tracts, in which stacked C:C+ bases increase the global pKa and stabilize the resulting iM^[7] [23]^. Our results show a positive correlation between increasing C-tract length and SNR across all conditions, indicating a role in iM stabilization, although at pH 5.5 and 40% BSA, this trend was dampened. Telomeric and centromeric iMs, which generally have tracts of three cytosines^[24]^, displayed a fairly consistent signal distribution. Negative control sequences did not yield a measurable response, regardless of screening condition, validating the specificity of the iMab antibody. Although there are examples in the literature of both stable Class I iMs with short loops (*VEGF*), and Class II iMs with long loops (*c-MYC*), we observe that shorter loops (*n* = 1-4), with thymine residues directly flanking C-tracts, are preferred for enhanced *in vitro* iM stability.

**Figure 1.**
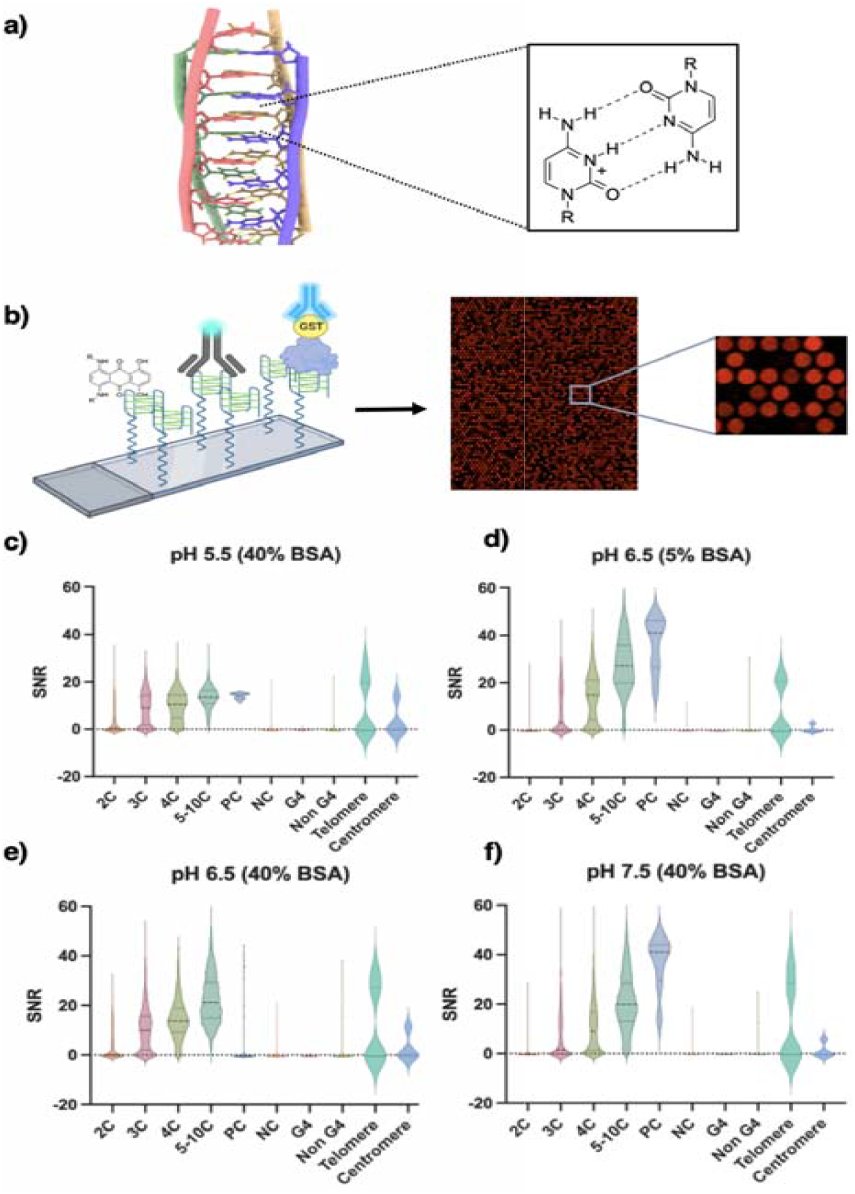
Using custom DNA microarrays to interrogate iM binding events a) Atomic resolution structure of an iM (PDB: 2N89[1] [10]) and illustration of the corresponding C:C+ basepair b) Illustration of the general microarray screening workflow in which diverse fluorescent ligands are incubated with the array and scanned with a fluorescence imager to quantify individual binding events. c-f) Violin plots demonstrating impact of pH (5.5 to 7.5) and molecular crowding (5% or 40% BSA) on iMab binding to diverse iM-forming and control sequence classes on the array

Select array iMs were analyzed to elucidate sequence-specificities that contribute to variations in iM folding or iMab binding by circular dichroism (CD) spectroscopy. iM 1 and 2, found in human genes *PELI3* and *SNX29* respectively, were chosen because they yielded high and low SNRs upon iMab incubation, despite having similar sequences (Figure 2a). iM 3, found in the *KIAA1614* gene, was chosen because it contains a flexible hairpin within its first loop, possibly conferring additional stability or influencing its binding profile^[25]^. Conversely, iM 4 is a synthetic variant of iM 3 in which select adenines were changed to thymines to abolish the hairpin, allowing for the observation of structure-specific differences in stability. Lastly, iM 5, found in many human promoters including *MAPK15, GPBP1L1, CCNI2*, and *TMC2*, was used as a reference, as it has been indicated to form a stable iM in previous studies. Figure 2b illustrates the impact of increasing pH on the iM 5 sequence. iM 5 exhibits a characteristic positive ellipticity at 287 nm and a negative ellipticity at 264 nm. At pH 4.5, the iM exhibits a classic spectra indicative of a fully folded iM structure (Figure 2b). As the pH is increased to 6.5, a hypochromic shift in the peaks is observed, indicating a transition from a stable iM to its partially state. When the solution becomes basic (pH 8), the iM is unraveled, yielding predominantly random coils. This was expected as iM 5 contains 4C-tracts which increases the global pKa of the ensemble through stacking interactions between C:C+ bases.^[26]^ Although there is conflicting evidence about the influence of loop length on iM stability, these results suggest that shorter loops are beneficial for increased stability and hydrogen bonding. CD melting experiments at 287 nm were performed at pH 4.5 and pH 6.5 (Figure 2a, 2c). Less acidic conditions decreased the melting temperature of all five iM sequences significantly, further demonstrating the pH dependence of iMs. At pH 4.5, the positive control (iM 5) exhibited the highest melt temperature (T_M_) of 60.9 ± 6.7 °C. iM 1, which exhibited the highest SNR upon iMab recognition, had the second highest T_M_ of 59.3 ± 7.3 °C. iM 2, despite its weak interaction with the iMab antibody, had a T_M_ of 58.8 ± 4.9 °C. We observe sequence-specific hypochromicity at 287 nm, with iM 1 and iM 2 which exhibited the highest and lowest CD absorption value, consistent with the SNR values reported in our iMab screen. Thus, while iMab recognizes diverse iM-forming sequences, it may have reduced recognition of iMs that are partially unfolded or contain GC-rich loops. Contrasting iM 3 and 4, we observe that abolishing the loop hairpin only confers a marginal increase in thermostability in some cases. At pH 4.5, the melting temperatures of iM 3 and 4 were similar at 46.1 °C and 47.0 °C respectively. At pH 6.5, however, iM 3 had a higher melting temperature than iM 4, indicating that hairpin may provide additional stabilization near physiological pH. We next sought to understand how sequence variations in iMs impact the recognition of a small molecule therapeutic. Mitoxantrone is reported to stabilize or destabilize various telomeric iM structures, and is used in the treatment of leukemia, breast cancer, and Non-Hodgkin’s lymphoma^[27]^. Contrary to iMab, which shows a strong positive correlation between SNR and iM C-tract length, mitoxantrone binding trends are less well-defined. With positive control iM sequences, we observe a dose-dependent increase in SNR (Figure 3d), however mitoxantrone also binds much more promiscuously to unstructured negative control sequences (Figure 3e). Additionally, mitoxantrone bound to various G4 array sequences. This is likely due to mitoxantrone’s planar structure, in which the anthraquinone core can stack or intercalate between nucleotides, allowing the secondary amines to interact electrostatically with phosphate backbones^[28]^. This nonspecific binding mode most likely mitigates mitoxantrone’s selective recognition of genomic iMs.

**Figure 2.**
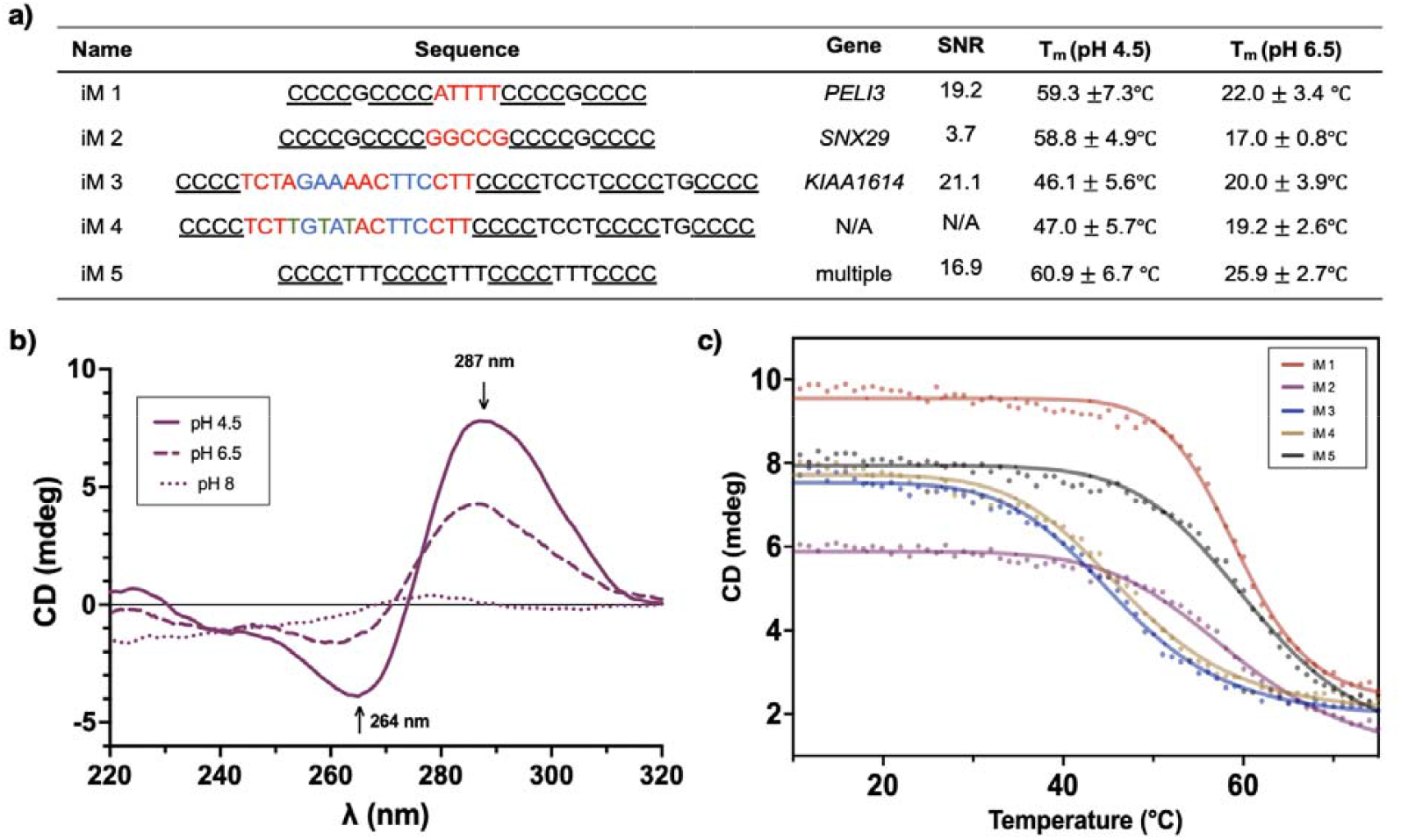
Circular dichroism spectroscopy and thermal melting assays of select iM-forming sequences. a) Table containing the array iM sequences, their genomic loci, normalized binding SNR from array, and melt temperature. b) CD spectra of iM 5 at pH 4.5, 6.5, and 8 c) Thermal melting curves of iM 1-5

**Figure 3.**
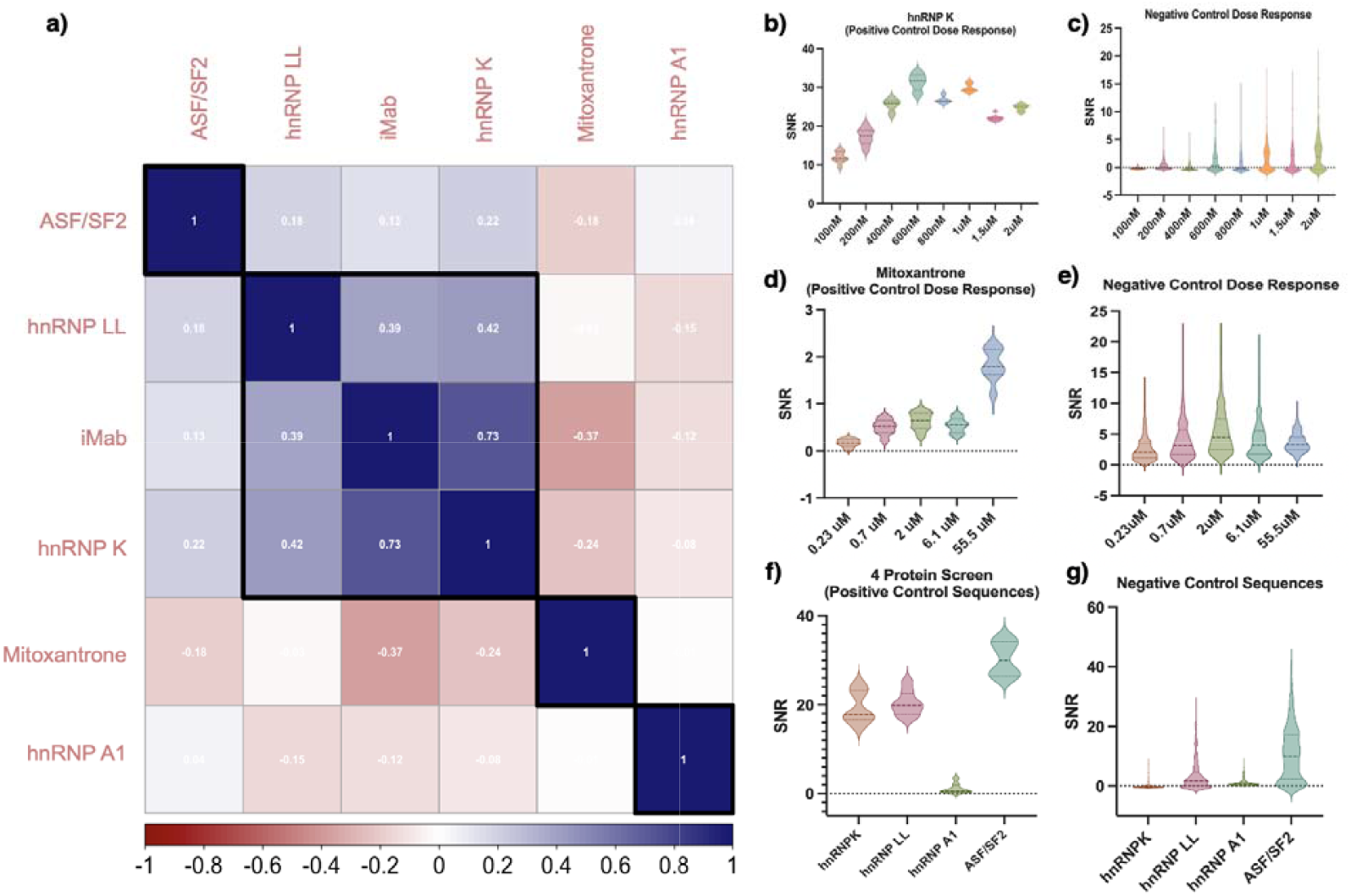
Analysis of protein and small molecule binding to iM sequences. a) Pearson correlation matrix comparing iM microarray screens. b) Violin plots illustrating median SNR distributions for a dose-response of hnRNP K with positive control array iMs and c) negative control sequences. d) Dose-response of mitoxantrone against positive controls and e) negative controls. f) Median SNR values for hnRNP K, hnRNP LL, hnRNP A1, and ASF/SF2 at 1.5 μM against positive controls and g) negative controls

We then examined four endogenous transcription factors that play important roles in RNA processing, transcriptional regulation, and genome maintenance-hnRNP K, hnRNP LL, ASF/SF2, and hnRNP A1. hnRNP K and hnRNP LL are members of the poly-(C) binding protein (PCBP) family, and have been shown to bind the *BCL2* iM, and the *KRAS* and *c-MYC* iMs, respectively. ^[29][30]^ These proteins bind to the lateral loops of the iM, partially unwinding the assembly to permit transcription. Dysregulation of hnRNP K is implicated with progression of breast cancer, lung cancer, prostate cancer, and acute myeloid leukemia^[31]^. hnRNP A1 recognizes G/C elements found in iM structures upstream of transcription start sites in *KRAS* and *HRAS*^*[32]*^. However, hnRNP A1 has also been shown to bind G4s within the 5’ UTR and promoters of *KRAS*^*[33]*^, *TRA2B*^*[34]*^, and *RON*, in a mechanism that is associated with metastatic relapse^[35]^. We expected the binding profile of hnRNP A1 to be skewed in favor of G-rich sequences in contrast to the other hnRNP proteins. ASF/SF2 is an oncoprotein transcription factor that interacts with hnRNP A1 to promote pre-mRNA splicing, and binds some of the same iMs as hnRNP K^[19]^. To elucidate differences between each protein binding profile, an all-against-all correlation analysis was performed (Figure 3a). The iMab antibody and hnRNP K binding profiles are positively correlated with each other (0.73), suggesting a strong preference for stable iM structures. hnRNP LL exhibited a weaker correlation, but still met the threshold to be clustered with iMab (0.39) and hnRNP K (0.42). Mitoxantrone and hnRNP A1 did not exhibit a positive correlation with any other protein that was screened. ASF/SF2 exhibited weak positive correlations with all proteins screened except mitoxantrone (-0.18). Our results suggest that hnRNP K is the most selective iM-binding protein of those that were screened. hnRNP K bound iM sequences in a dose-dependent manner that saturated around 600 nM (Figure 3b). hnRNP K, hnRNP LL, and ASF/SF2 tightly bound to positive control iM sequences, with ASF/SF2 exhibiting the highest median SNR. In contrast, hnRNP A1 did not yield a significant SNR distribution for any positive control iMs. hnRNP K and hnRNP A1 did not bind to negative control sequences on the array, whereas ASF/SF2 and hnRNP LL bound some negative controls, suggesting broad nucleic acid recognition beyond iMs (Figure 3f, 3g). We visualized all normalized array interactions in a heatmap delineated by sequence class (Figure 4a). hnRNP LL bound considerably more G4 and negative control sequences than hnRNP K, despite them both being PCBPs. The RRM1 iM-recognition motif of hnRNP LL, and its interaction with the *BCL2* promoter iM has been well characterized^[36]^, and is supported by evidence that it binds (CA)_*n*_ dinucleotide repeats in 3’ UTR regions of pre-mRNA^[37]^. However, hnRNP LL also represses transcription via the *Gad1* promoter G4 in neurons when synaptic activity is high^[38]^. Such competitive interactions could explain its reduced affinity for iMs and off-target recognition compared to hnRNP K. hnRNP A1 is an early tumor biomarker, and works with ASF/SF2 in antagonistic fashion to regulate alternative splicing, such as in the case of *IRF-3*^*[39] [40]*^. hnRNP A1 is reported to recognize G/C sequence elements and its interactions with neuronal, telomeric, and oncogenic G4s are well documented^[33, 41]^. Likewise, hnRNP A1 exhibited a strong preference for DNA G4s on the array. Its binding profile was plotted against hnRNP K to examine divergences in binding preferences by sequence class (Figure 4b). It is evident that hnRNP K exhibits a clear preference for iMs with a minimum of three cytosines in the repeat. hnRNP A1 exhibits a broad binding profile, preferentially binding to less-stable iMs with short cytosine tracts, unstructured negative control sequences, and G-quadruplexes. Over all sequence classes represented on the array, hnRNP K and hnRNP A1 exhibit very little, if any correlation. It has been proposed that these proteins belong to distinct subgroups within the hnRNP family, where hnRNP A1 is involved in core complex formation and hnRNP K plays roles in RNA regulation^[42]^. Recently, it was shown that hnRNP A1, R, U, and K form a complex in colon and rectal adenocarcinoma cells, in which hnRNP K regulated translation of relevant genes^[43]^. When cross-referencing the published co-expression network of the hnRNP A1/R/U/K complex with the array hnRNP K data, we see that several genes identified are hits. These hits were found to be the 4C iMs *TARDBP* (SNR: 26.87), *MRPS24* (SNR: 25.64), and *RAD23B* (SNR: 8.21). hnRNP A1 may locally encourage hnRNP K iM binding through complex formation and subsequent conformational shifts. The interaction between hnRNP K and A1 to maintain cellular homeostasis, and the associated role that iM recognition plays, warrants further study to shed light on potential molecular targets. Lastly, ASF/SF2 bound the most stable iM structures on the array, sequences with extended C-tracts however, like hnRNP A1, ASF/SF2 bound to a majority of the G4 sequences on the array (Figure 4c). Indeed, ASF/SF2 recognizes G4-forming r(GGGGCC)_*n*_ repeats found in UTRs and intronic regions of pre-mRNA^[44]^, such as in *C9orf72*, which is also bound by hnRNP A1^[45]^, explaining these observations. Looking further into the hnRNP K iM binding profile, we observe clear correlations between hnRNP K binding propensity, C-tract length, loop length, and loop GC content. More specifically, iM loop regions between one to four nucleotides long are ideal for hnRNP K recognition for all three loop regions in the array iMs (Figure 5b). We see a negative correlation between increasing loop GC content and iM stability, which further confirms the findings that loop regions with adenines or thymines confer additional stability through hydrogen bonding (Figure 5c). The *c-MYC* promoter iM, which is recognized by hnRNP K’s KH domains, contains 5C-tracts and four loops of one adenine or thymine^[46]^. We subsequently set out to determine if these results correlated with hnRNP K’s regulatory targets *in vivo*. We compared our *in vitro* array data to ChIP-Seq data from an RNA-binding protein interactome dataset from ENCODE. We used data from two different human cell lines, HepG2, a liver cancer cell line, and K562, a myelogenous leukemia cell line (see Methods)^[47]^. The SNR values from the arrays show positive correspondence (R = 0.12 to 0.34, depending on iM class) with maximum ChIP-Seq peak density in the HepG2 cell line (Figure 5d), highlighting agreement of our array-based measurements with *in vivo* binding data. This correlation is lower for the K562 cell line (R = 0.053 to 0.21), suggesting that additional cell-specific factors impact hnRNP K binding *in vivo*. Furthermore, we find that the fraction of significantly enriched (P_adj_ < 0.05) iM sequences in ChIP-Seq peaks increases with SNR (Figure 5e). Specifically, 35% (31/88) and 25% (22/28) of highly bound (SNR > 40) iM array sequences are enriched in hNRNP K ChIP-Seq peaks from HepG2 and K562 cells respectively, suggesting that the binding preferences observed on the arrays are reflected *in vivo*. This suggests that array sequences with a high SNR show a positive correlation with endogenous nucleic acid targets of hnRNP K in two human cancer cell lines.

**Figure 4.**
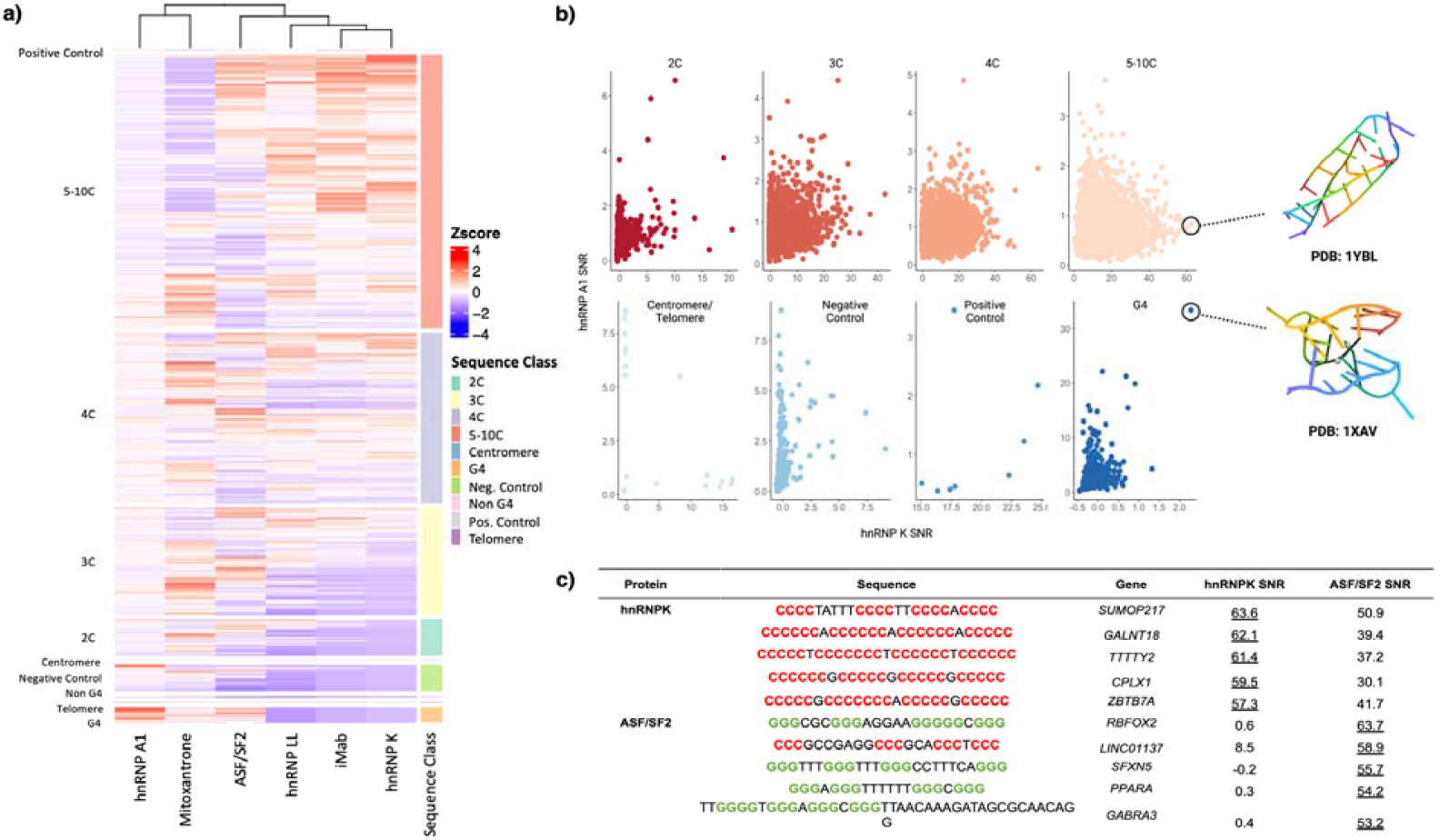
Comparison of iM binding profiles. a) Hierarchically clustered heatmap depicting normalized SNR values for all screens. Heatmap is arranged by the sequence classes represented on the array. b) Scatterplots contrasting hnRNP K binding profile wit hnRNP A1 binding, highlighting class specific binding differences. The color of the points is scaled by hnRNP K SNR. c) Sequence table comparing top hits bound by hnRNP K and ASF/SF2

**Figure 5.**
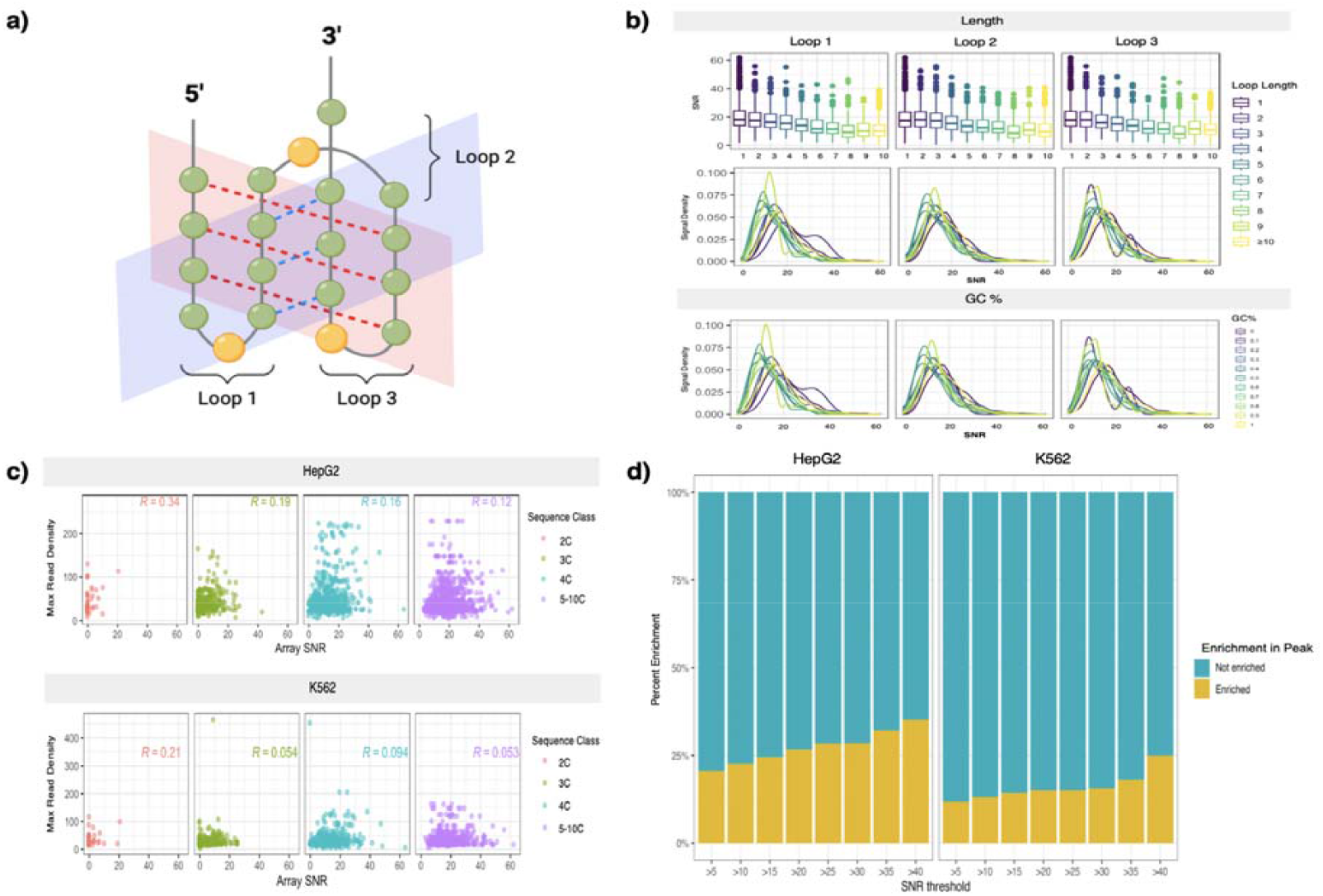
hnRNP K binds to stable iMs in biochemical and cellular assay. a) Graphical representation of iM structure and loop regions, where the green circles represent cytosines and yellow circles represent any loop residue b) Analysis of the impact of iM loop length on hnRNP K recognition. Box plots and signal density plots for hnRNP K binding to 5-10 C-containing iMs of increasing loop length (from 1 to 10 nucleotides) and GC content (from 0 to 100%). (c) Correlation analysis between hnRNP K binding to iM sequences on arrays and in cells. Plot axes compare maximum ChIP-Seq read density for hnRNP K peaks in HepG2 and K562 cells (from ENCODE) to iM array SNR for HepG2 and K562 cells. d) Fraction of tightly-bound array sequences (top 10%) represented in hnRNP K ChIP-Seq peaks in HepG2 and K562 cell lines at different iM array SNR thresholds.

## Conclusion

In summary, we demonstrate that microarrays can quantiatively assess thousands of interactions between various biomolecules, and genomic iM sequences, revealing both sequence and structure-specific binding information. We establish that iM formation is influenced by pH, C-tract length, loop length, loop GC%, and molecular crowding, and determine optimal conditions for their stable formation on the array. Based on these constraints, we identify orthogonal iM sequence and structure-based binding preferences exhibited by iMab, mitoxantrone, hnRNP K, hnRNP LL, hnRNP A1, and ASF/SF2. We observe that iMab readily binds Class I iMs (with shorter loops) in acidic solutions, and Class II iMs (with longer loops) near physiological pH in the presence of crowding. iM loop GC% also impacts recognition, in which lower loop GC% is favorable. These observations reinforce the utility of iMab as a powerful tool for the recognition of diverse iM sequences from the genome. These trends are mirrored by hnRNP K, which exhibits a preference for genomic iMs with 3-5 cytosine repeats, loop regions between 1-3 nucleotides, and loop GC% between 0-20%. Array-based findings were cross-referenced against open-access ENCODE ChIP-Seq data, where well-bound array sequences positively correlated with hnRNP K ChIP-Seq read density in two human cancer cell lines. Taken together these data provide important insights into the fine-tuned recognition of iM structures by multiple disease-relevant transcription factors and may help inform the rational design of therapeutics that target such interactions.

## Supporting Information

Experimental procedures, computational methods, and code/data availability are included in the Supporting Information.

## Acknowledgements

This research was supported by the Intramural Research Program of the National Institutes of Health, National Cancer Institute, Center for Cancer Research, project number Z01 BC011585 07 (PI, J. S. Schneekloth, Jr). This work utilized the computational resources of the NIH HPC Biowulf cluster (http://hpc.nih.gov). Additionally, we thank Dr. Sergey Tarasov and Marzena Dyba of the National Cancer Institute Biophysics Resource for allowing us to use their instrumentation and appropriately interpret biophysical data.

