## Supplemental Information for "DECODING COMPLEXITY IN BIOMOLECULAR RECOGNITION OF DNA I-MOTIFS"

**DECODING COMPLEXITY IN BIOMOLECULAR RECOGNITION OF DNA I-MOTIFS WITH MICROARRAYS**

**Abstract:** DNA i-motifs (iMs) are non-canonical C-rich secondary structures implicated in numerous cellular processes. Though iMs exist throughout the genome, our understanding of iM recognition by proteins or small molecules is limited to a few examples. We designed a DNA microarray containing 10,976 genomic iM sequences to examine the binding profiles of four iM-binding proteins, mitoxantrone, and the iMab antibody. iMab microarray screens demonstrated that pH 6.5, 5% BSA buffer was optimal, and fluorescence was correlated with iM C-tract length. hnRNP K broadly recognizes diverse iM sequences, favoring 3-5 cytosine repeats flanked by thymine-rich loops of 1-3 nucleotides. Array binding mirrored public ChIP-Seq datasets, in which 35% of well-bound array iMs are enriched in hnRNP K peaks. In contrast, other reported iM-binding proteins had weaker binding or preferred G-quadruplex (G4) sequences instead. Mitoxantrone broadly binds both shorter iMs and G4s, consistent with an intercalation mechanism. These results suggest that hnRNP K may play a role in iM-mediated regulation of gene expression *in vivo*, whereas hnRNP A1 and ASF/SF2 are possibly more selective in their binding preferences. This powerful approach represents the most comprehensive investigation of how biomolecules selectively recognize genomic iMs to date.

DOI: 10.1002/anie.2021XXXXX

Table of Contents

Experimental Section and Computational Methods	2

SOURCES OF ANTIBODY, SMALL MOLECULE, AND RECOMBINANT PROTEIN	2

MICROARRAY BINDING ASSAYS	3

SPECTRAL AND THERMAL MELTING ANALYSIS BY CIRCULAR DICHROISM	3

DATA PROCESSING AND ANALYSIS	3

CORRELATION ANALYSIS WITH ENCODE ChIP-Seq DATA	3

DATA AVAILABILITY	4

Results and Discussion	4

Figure S1. Circular dichroism (CD) spectra of i-Motif 1 (iM 1) at pH 4.5, 6.5, and 8	4

Figure S2. Circular dichroism (CD) spectra of iM 2 at pH 4.5, 6.5, and 8	4

5

Figure S3. Circular dichroism (CD) spectra of iM 3 at pH 4.5, 6.5, and 8	5

Figure S4. Circular dichroism (CD) spectra of iM 4 at pH 4.5, 6.5, and 8	5

Figure S5. Hierachichally clustered pairwise correlations for all DNA iM microarray screening conditions performed. Pearson R correlation coefficients are displayed and scaled by color.	6

Figure S6. Signal distributions for all DNA iM microarray screening conditions separated by sequence class.	8

Figure S7. Heatmap depicting raw SNR values on the array with increasing hnRNP K concentration. Sequence class is shown on the right with the corresponding color shown.	9

Figure S8. Impact of iM loop length, from one to ten, on protein binding. The color corresponding to each iM loop length is shown on the right.	9

Figure S8. Impact of iM loop GC%, from 0% to 100%, on protein binding. The color corresponding to each iM loop GC% is shown on the right	10

Figure S9. The the correlation between hnRNP K array SNR and max read density from hnRNP K peaks in public ChIP-Seq data from both HepG2 and K562.	11

Experimental Section and Computational Methods

SOURCES OF ANTIBODY, SMALL MOLECULE, AND RECOMBINANT PROTEIN

Mitoxantrone was purchased from Ambeed (1,4-Dihydroxy-5,8-bis((2-((2-hydroxyethyl)amino)ethyl)amino)anthracene-9,10-dione, Cat. No. A207951). We obtained the anti-i-motif DNA antibody conjugated with FluoProbes®647H (Cy5-BG4) from Absolute Antibody (product number Ab01462-23.0). N-terminal Glutathione S-transferase (GST) tagged human ASF/SF2, hnRNP K, hnRNP LL hnRNP A1 were synthesized by GenScript.

MICROARRAY BINDING ASSAYS

Microarrays were pre-equilibrated in a PBS buffer containing 100 mM KCl, 0.01% Triton X-100 buffer for 1 hour at room temperature. They were subsequently blocked with the same buffer containing 2% nonfat dry milk, and washed without detergent, to properly fold array sequences. For all screens, a binding solution of PBS, 100 mM KCl, 51.3 ng/µL salmon testes DNA, a varied concentration of BSA (%v/v), and an appropriate concentration of GST-tagged protein, antibody, or small molecule was added. For certain experiments the pH of the buffer was also varied. Microarrays were incubated for 1 hour at room temperature, and washed with DI H_2_O. For GST-tagged protein screens, the slide was incubated in a binding solution containing an anti-GST antibody, and the process was repeated as described above.

SPECTRAL AND THERMAL MELTING ANALYSIS BY CIRCULAR DICHROISM

Spectral analysis was performed using a J-1500 CD spectrophotometer and 1.0 cm path-length quartz cells at 25 °C. iM DNA sequences were annealed at 95℃ for 2 minutes, in 2X PBS, 100 mM KCl buffer at varying pHs. Each spectral measurement was taken in triplicate to obtain an average spectra from 220-340 nm. For thermal melting assays, iM DNA sequences were folded in 2X PBS, 100 mM KCl pH 4.5 buffer at 5 µM at 95℃ for 2 minutes. 300 µL of DNA solution was added to a 10mm quartz cuvette, and heated from 20 to 95℃ in a CD spectrometer (JASCO 1500) with a temperature step interval of 1 ℃/min. The melting temperature (T_m_) was determined by continuously measuring the absorbance at 287 nm across the temperature gradient. The temperature-dependent response was plotted, and the T_m_ value was determined as the peak of the first derivative curve using GraphPad Prism 8 software.

DATA PROCESSING AND ANALYSIS

All 8x60K Agilent microarray slides were scanned using an Innopsys InnoScan 1100 using extended dynamic range (XDR), 0.5 µm pixel size, and the 635 nm scan setting. SNR spot intensities from microarray TIF images were extracted using the Agilent Feature Extraction Software and are reported as raw fluorescence units. For each unique probe on the array, the median SNR was computed across all probes containing an identical probe identifier.

CORRELATION ANALYSIS WITH ENCODE ChIP-Seq DATA

We downloaded human hnRNP K ChIP-Seq peak coordinates for K562 (accession: ENCFF505RNR) and HepG2 (accession: ENCFF035OPG) cell lines from the ENCODE website (https://www.encodeproject.org/)^[48-49]^. For each iM array sequence, we computed an enrichment score, $E=\frac{\mathrm{OCC}_{\mathrm{obs}}}{\mathrm{OCC}_{\exp}}$, within ChIP-Seq peaks in each cell line^[50]^. *OCC_obs_* is the observed occurrence of each iM sequence in the set of ChIP-Seq peaks, and OCC_exp_ is the expected occurrence given the frequency of the iM sequence across the human genome, computed as $\mathrm{OCC}_{\exp}=N\times\frac{L_{r}}{L_{g}} ,$ where *N* is the total number of motifs in the genome, *L*_r_ is the total length (in base pairs) in the subset of peaks, and *L_g_* is the total length (in base pairs) of the human genome. For each iM sequence, a p-value was computed using a chi-squared test, and adjusted for the false discovery rate using the Benjamini-Hochberg method. Additionally for each iM sequence on the array, we computed an *in vivo* binding score, which is the maximum read density observed across all ChIP-Seq peaks containing the probe sequence. Code is available at [https://github.com/desireetillo/imotif2023](https://gcc02.safelinks.protection.outlook.com/?url=https%3A%2F%2Fgithub.com%2Fdesireetillo%2Fimotif2023&data=05%7C01%7Cjay.schneekloth%40nih.gov%7Cfd0a88c49e904c61c16408db320f3caa%7C14b77578977342d58507251ca2dc2b06%7C0%7C0%7C638158813784005824%7CUnknown%7CTWFpbGZsb3d8eyJWIjoiMC4wLjAwMDAiLCJQIjoiV2luMzIiLCJBTiI6Ik1haWwiLCJXVCI6Mn0%3D%7C3000%7C%7C%7C&sdata=YThgJF6E404au7uEFCao%2BbYH%2BwPchsBSnrokPXt1n%2Fc%3D&reserved=0).

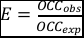

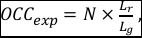

DATA AVAILABILITY

Data (array probe sequences, raw probe intensities, and median feature intensities) are available at the NCBI GEO database under accession #GSE227616).

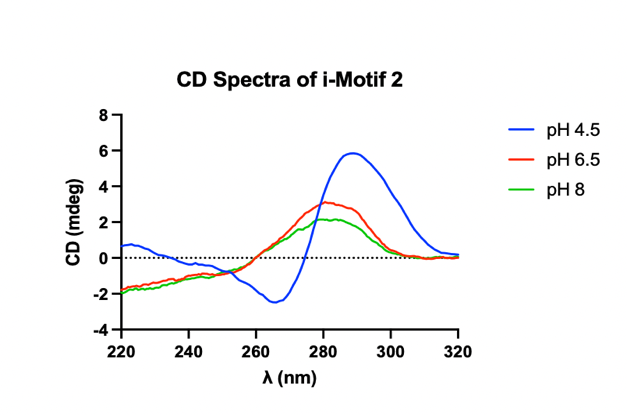

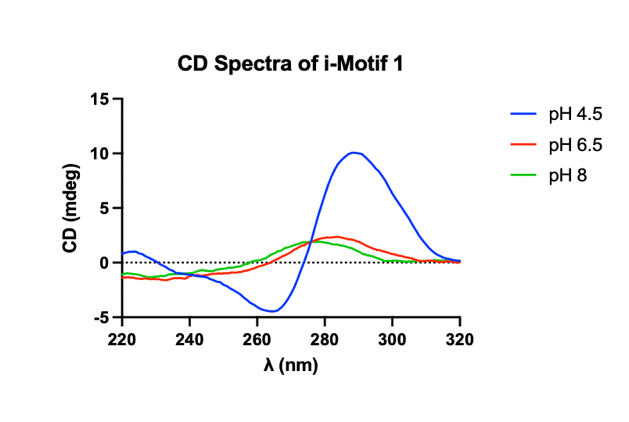
Results and Discussion

Figure S1. Circular dichroism (CD) spectra of i-Motif 1 (iM 1) at pH 4.5, 6.5, and 8

Figure S2. Circular dichroism (CD) spectra of iM 2 at pH 4.5, 6.5, and 8

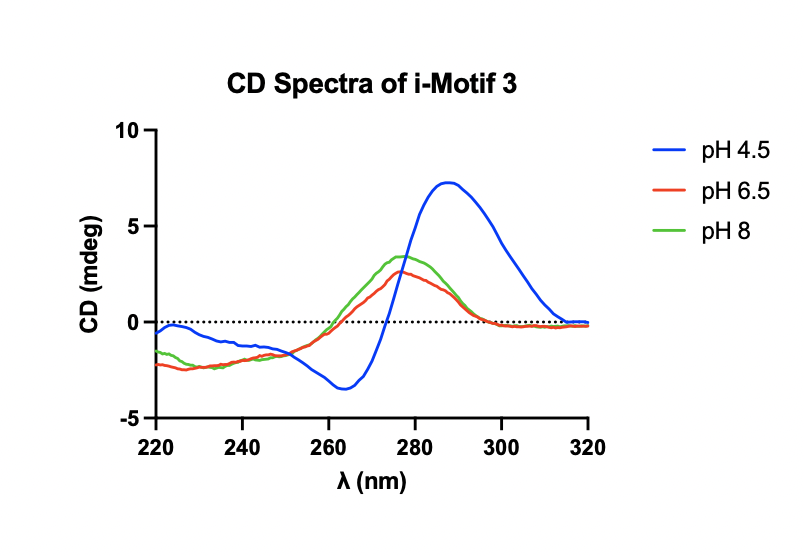

Figure S3. Circular dichroism (CD) spectra of iM 3 at pH 4.5, 6.5, and 8

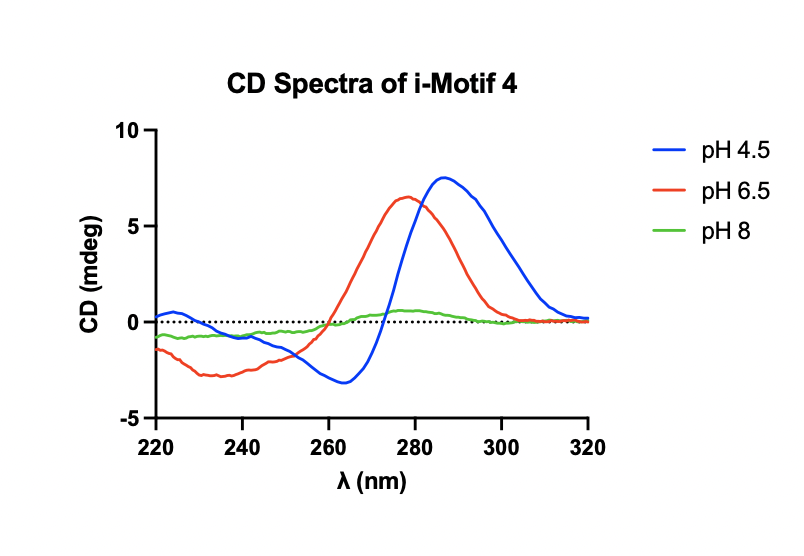

Figure S4. Circular dichroism (CD) spectra of iM 4 at pH 4.5, 6.5, and 8

Figure S5. Hierachichally clustered pairwise correlations for all DNA iM microarray screening conditions performed. Pearson R correlation coefficients are displayed and scaled by color.

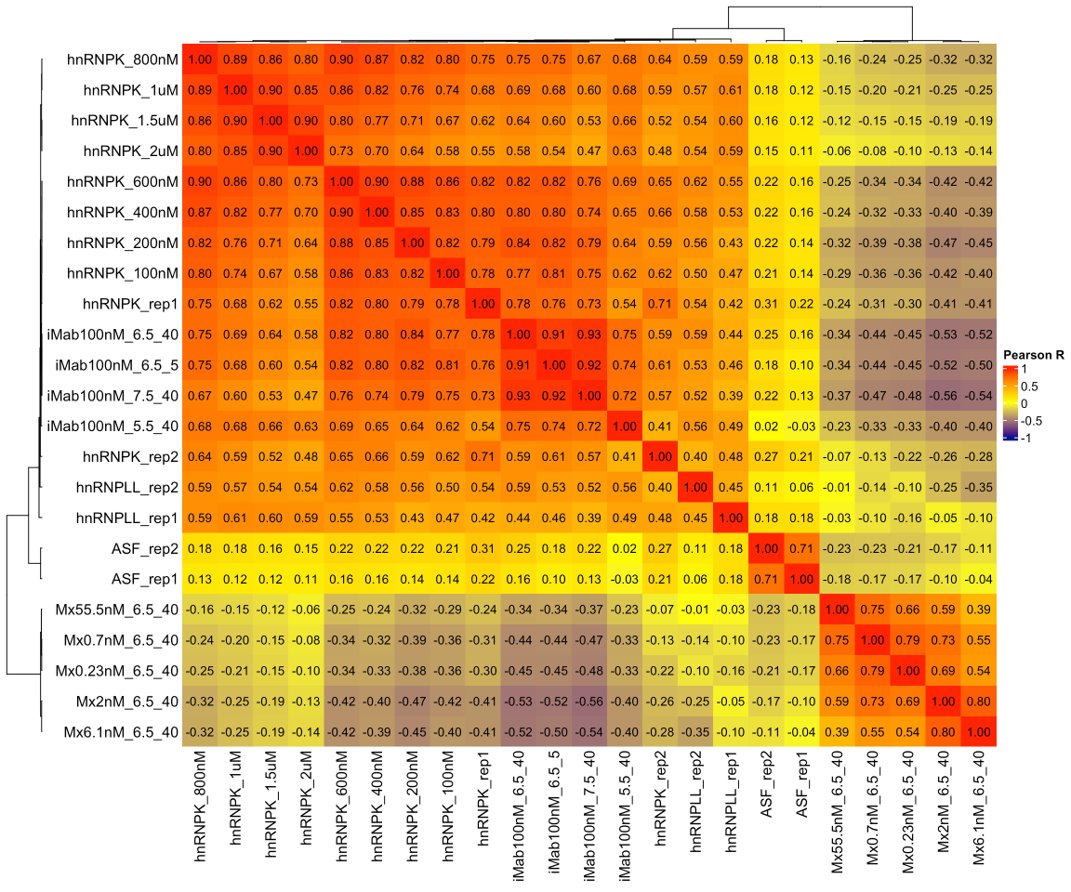

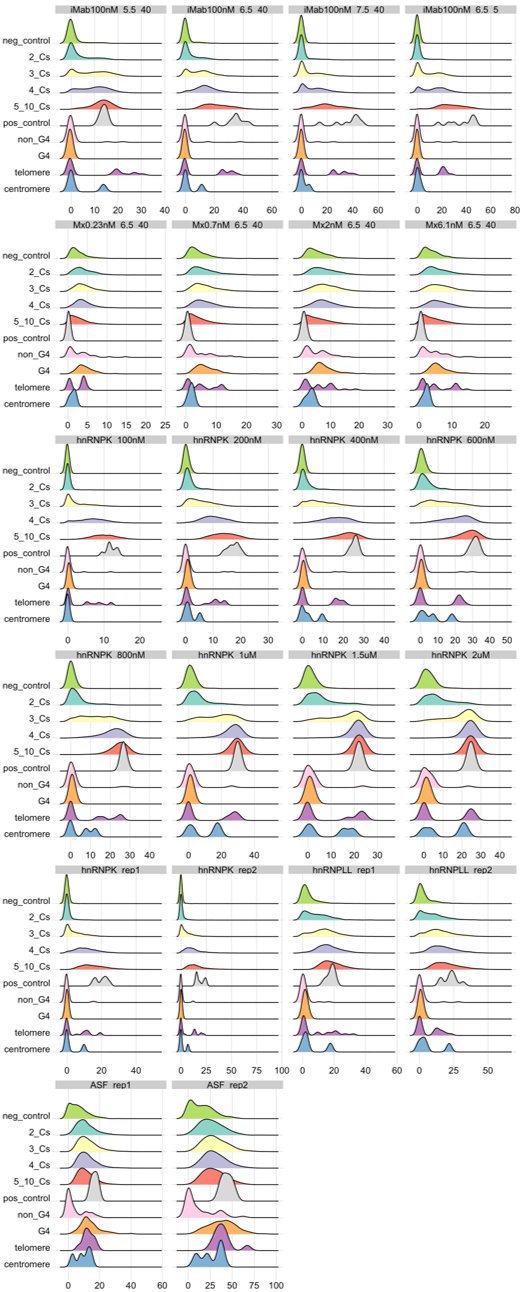

Figure S6. Signal distributions for all DNA iM microarray screening conditions separated by sequence class.

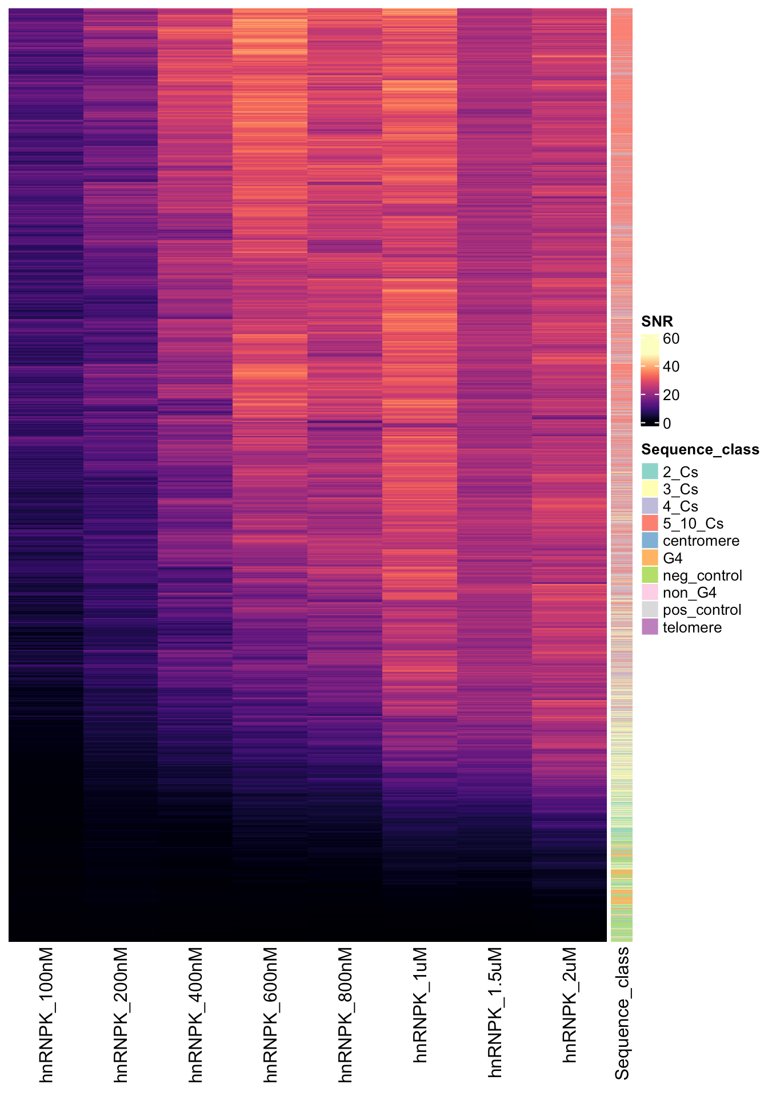

Figure S7. Heatmap depicting raw SNR values on the array with increasing hnRNP K concentration. Sequence class is shown on the right with the corresponding color shown.

Figure S8. Impact of iM loop length, from one to ten, on protein binding. The color corresponding to each iM loop length is shown on the right.

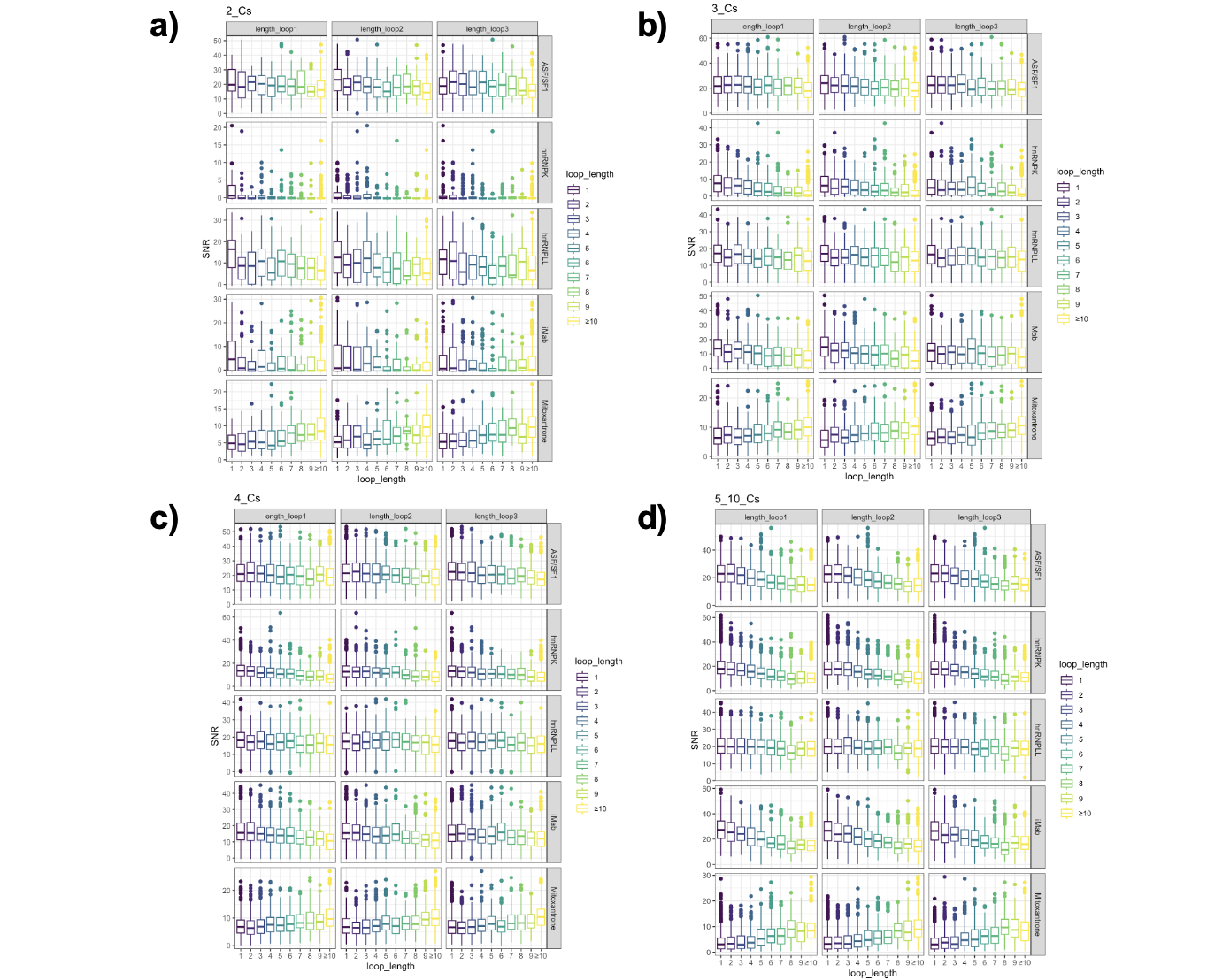

Figure S8. Impact of iM loop GC%, from 0% to 100%, on protein binding. The color corresponding to each iM loop GC% is shown on the right

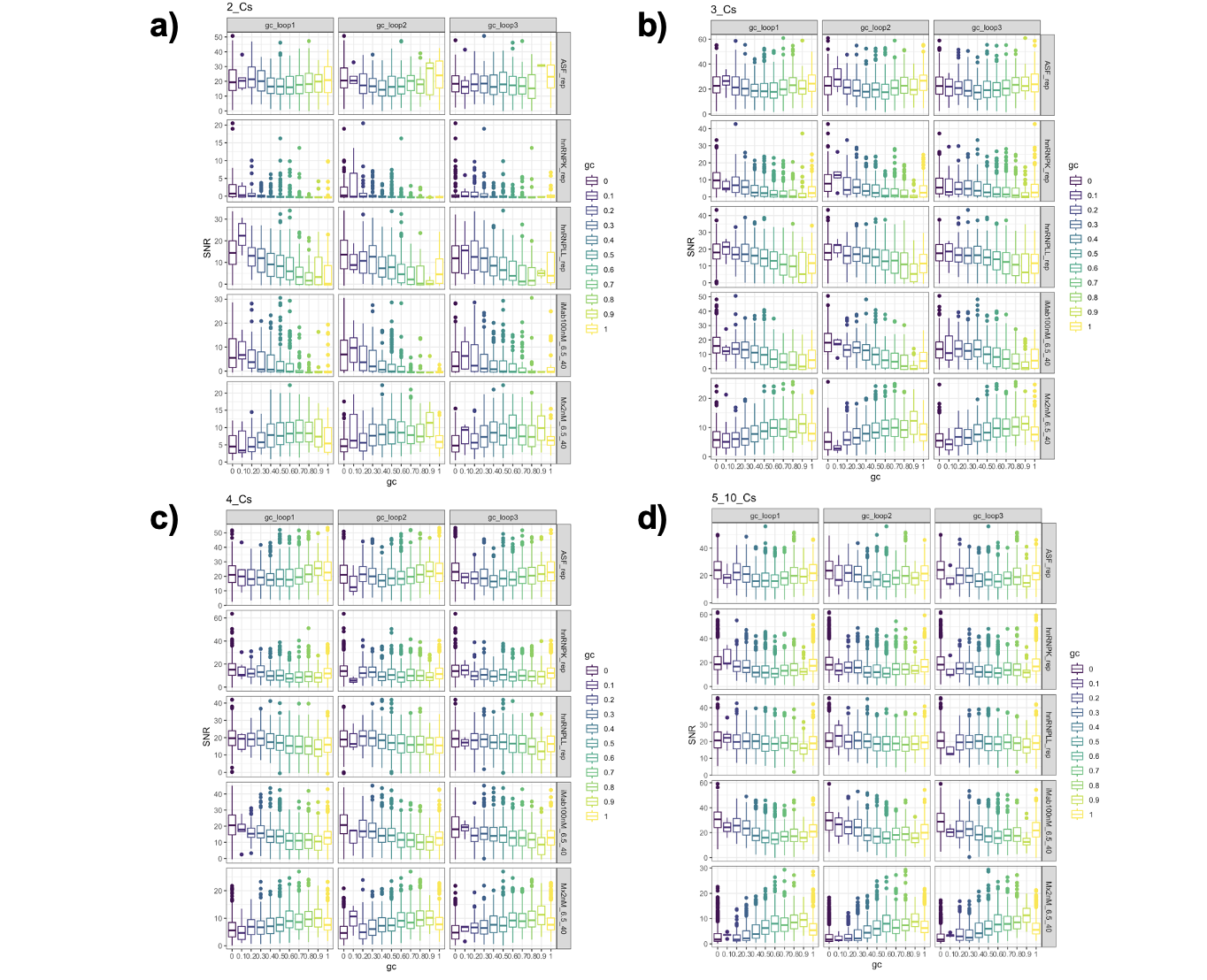

Figure S9. The the correlation between hnRNP K array SNR and max read density from hnRNP K peaks in public ChIP-Seq data from both HepG2 and K562.
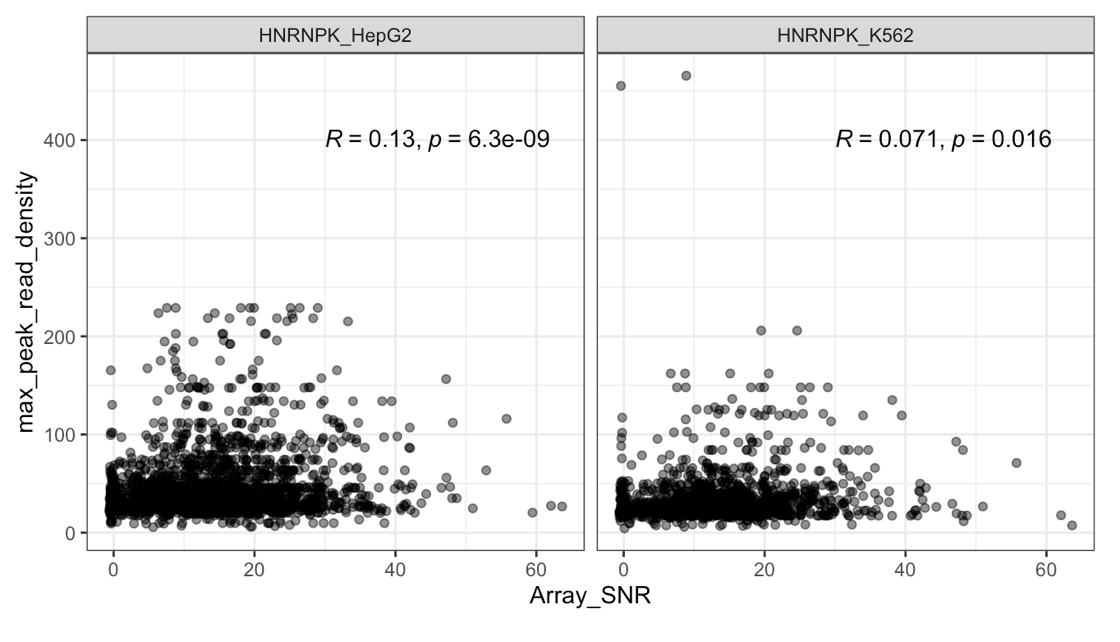
